# Cancer-Induced Immunosuppression can enable Effectiveness of Immunotherapy through Bistability Generation: a mathematical and computational Examination

**DOI:** 10.1101/498741

**Authors:** Victor Garcia, Sebastian Bonhoeffer, Feng Fu

## Abstract

Cancer immunotherapies rely on how interactions between cancer and immune system cells are constituted. The more essential to the emergence of the dynamical behavior of cancer growth these are, the more effectively they may be used as mechanisms for interventions. Mathematical modeling can help unearth such connections, and help explain how they shape the dynamics of cancer growth. Here, we explored whether there exist simple, consistent properties of cancer-immune system interaction (CISI) models that might be harnessed to devise effective immunotherapy approaches. We did this for a family of three related models of increasing complexity. To this end, we developed a base model of CISI, which captures some essential features of the more complex models built on it. We find that the base model and its derivates can plausibly reproduce biological behavior that is consistent with the notion of an *immunological barrier*. This behavior is also in accord with situations in which the suppressive effects exerted by cancer cells on immune cells dominate their proliferative effects. Under these circumstances, the model family may display a pattern of *bistability*, where two distinct, stable states (a cancer-free, and a full-grown cancer state) are possible. Increasing the effectiveness of immune-caused cancer cell killing may remove the basis for bistability, and abruptly tip the dynamics of the system into a cancer-free state. Additionally, in combination with the administration of immune effector cells, modifications in cancer cell killing may be harnessed for immunotherapy without the need for resolving the bistability. We use these ideas to test immunotherapeutic interventions *in silico* in a stochastic version of the base model. This bistability-reliant approach to cancer interventions might offer advantages over those that comprise gradual declines in cancer cell numbers.

## INTRODUCTION

Mathematical modeling of cancer-immune system interactions can reveal the fundamental mechanisms that govern the dynamics of tumor growth [1, 2], and represent and important tool to devise and test new forms of immunotherapy *in silico* [3]. The modeling relies on the appropriate integration of how these different cell types affect one another [4–8]. Recent studies have uncovered a plethora of interactions by which cancer cells affect immune cells, and vice versa [2, 9]. For instance, cancer cells elicit immune responses by a variety of effector cells [9– 11]. These effector cells, in particular white blood cells, natural killer cells (NKs) and cytotoxic T lymphocytes (CTLs) can lyse cancer cells [12], inhibiting tumor growth or even eliminating microscopic tumors altogether — a process termed *immunosurveillance* [13, 14]. However, cancers have also been shown to be able to suppress the proliferation of effector cells, which typically target cancer cells with specific biochemical signatures [15, 16]. Cancer cells accrue mutations that, by changing these signatures, enable them to partially evade recognition [1, 17, 18]. Furthermore, cancers may actively down-regulate immune responses elicited against them [19–23], for example by recruiting the action of T regulatory cells [9, 24, 25], leading to *immunosupression*. A summary of these interactions shows that all combinations of stimulation and suppression on growth between cancer and immune cells may act simultaneously (see Figure 1A). These interactions direct the interplay between the cancer and the immune system. Thus, their integration into mathematical models can reveal how immunotherapeutic approaches may be employed with maximum efficiency. The main immunotherapy approaches today work by impairing mechanisms that allow cancers to evade immune-recognition or by the administration of effector cells to the host [9, 26] (see Figure 1B).

**FIG. 1.**
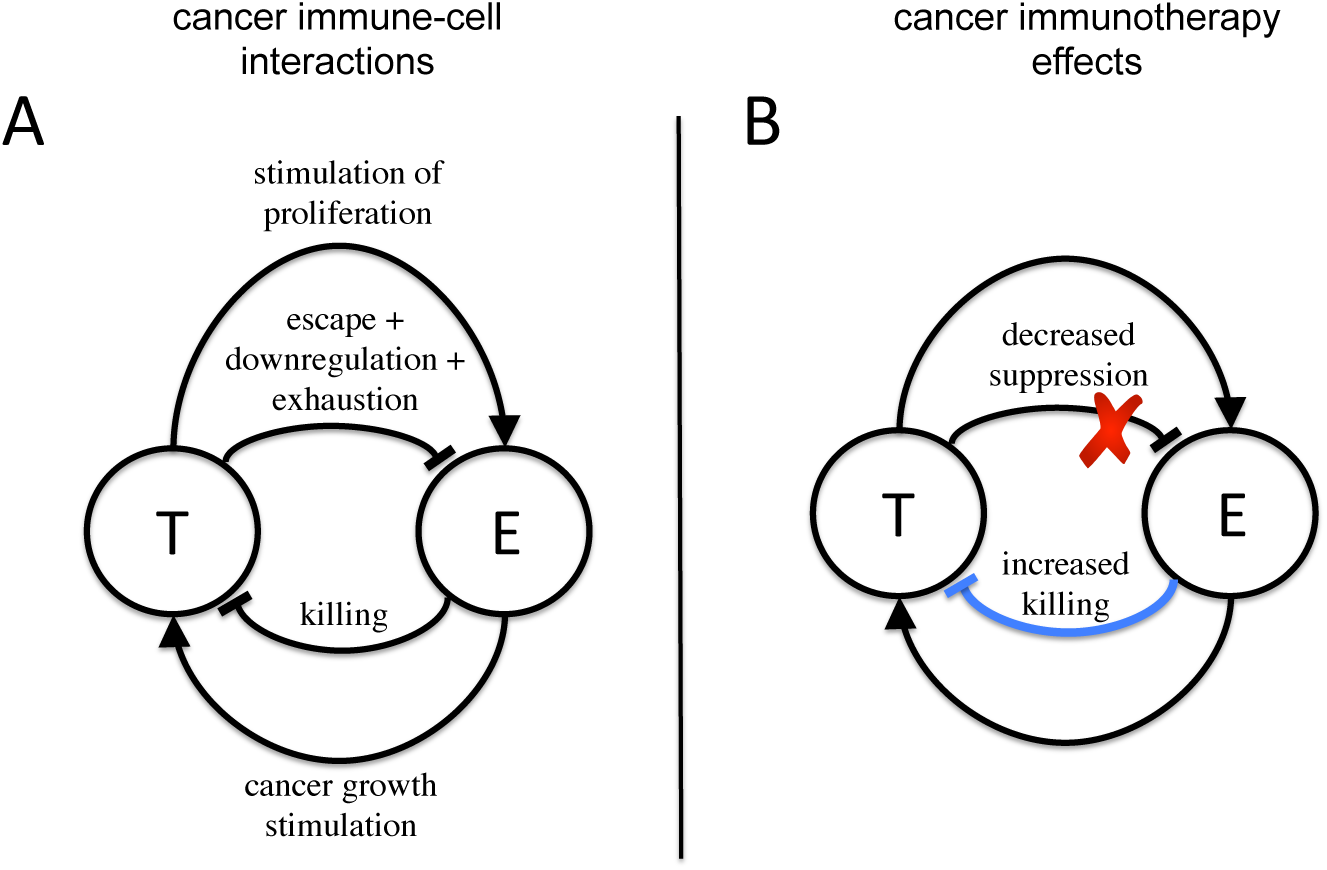
Cancer-immune system interactions and effects of immunotherapy. A) Interactions governing the dynamics between cancer cells (T) and immune cells targeting the cancer cells —termed effector cells— (E). A complex web of interactions has been identified [1, 2, 9], with both cell types capable to both stimulate and suppress each others’ growth. B) Immunotherapy acts by either increasing the killing rate of effector cells, for example by administrating new effector cells into the host (*adoptive T cell transfer*), or by impairing the escape mechanisms cancer cells adopt to avoid being cleared, for example by *monoclonal antibody therapy*.

Due to their importance for immunotherapy, mathematical models of cancer-immune interactions have been the focus of intense theoretical efforts over the last decade [1, 2]. At the heart of this theoretical effort lies the task of identifying what features of cancer-immune dynamics are most effectively used to achieve immunotherapeutic success, i.e. cancer eradication. Following this tradition, we focus on deterministic, population based non-spatial models for analysis. We do this for two reasons. First, their mathematical simplicity is better suited to unambiguously reveal those model properties that are crucial for the modification of a cancer’s state, since the number of factors influencing the dynamics is manageable and understandable [2]. Second, we can draw from a broad body of theory about immune system pressures developed within cancer research [2, 8, 27–31] as well as in virus dynamics, and in particular, human immunodeficiency virus (HIV) dynamics research [32–43]. To explore immunotherapeutic interventions *in silico*, we use a stochastic version of one of the models analyzed.

In this study, we have explored whether there exist simple, consistent properties of cancer-interaction models that might be harnessed to devise effective immunotherapy approaches. We investigated this question in a family of three related models of increasing complexity. To this end, we developed a base model of cancer immune system interaction (CISI), which captures some essential features of more complex models. The purpose of the model is thus to serve as a potentially useful guide for treatment therapies based on immune action. We analyzed under what parameter regimes the model produces biologically plausible behavior, and investigated how steady states are affected by changes in these parameters. We then successively extended the model to first include more complex features of cancer-immune system interactions [36], such as saturating proliferative stimulation and exhaustion and in a second step, to include the action of NKs and CTLs. The common properties identified in these models were then used to study how combinations of immunotherapeutic treatments may work together to achieve eradication. To this end, we implemented stochastic simulations of the base model and analyzed how the dynamics are affected by *adoptive T cell transfer* [44–47], as well as by the disruption of immune evasion mechanisms of the cancer through for example monoclonal antibody therapy [9] [48].

## MATERIALS AND METHODS

To analyze the algebraic properties of a system of equations involving cancer-immune interactions, we used the program *Mathematica* [49]. To find equilibrium points in situations where this was not algebraically possible, we used the *rootSolve* package in R [50–52].

Since all ordinary differential equations (ODEs) here described are deterministic, the time course of the decline of cancer cell numbers will always follow the same continuous trajectory given the identical initial conditions. However, when small cancer cell numbers are reached, the temporal order at which the discrete events occur that underpin the dynamics will become important. Such events include the replenishment of immune cells and cancer cell deaths. Thus, at small cell numbers, accounting for the stochasticity of these events will add realism to the simulation, and help decide when eradication has effectively been achieved. To this end, we employed the Gillespie algorithm, where the interactions between cell types are explicitly simulated. Stochastic simulations of all ODEs were run in the R language for statistical computing [53] by using the Gillespie algorithm [54] with tau leaping in the *adaptivetau* package [55]. If not stated otherwise, simulations were run with the set of parameter values given in Table I. For alternative strategies to account for the stochasticity of CISI at the temporal mesoscale see [56].

**TABLE I.** Standard parameter values for the base model.

| Parameter | Description | Value |
| --- | --- | --- |
| $a$ | maximum replication rate of cancer cells [5] | $0.514 \text{ day}^{-1}$ |
| $b$ | inverse carrying capacity of tumor [5] | $1.02 \cdot 10^{-9} \text{ cells}^{-1}$ |
| $k$ | killing efficiency of immune cells [57, 58] | $10^{-4} - 10 \text{ cells}^{-1} \text{ day}^{-1}$ |
| $\sigma$ | replenishment rate of immune cells [35] | $10 \text{ cells} \cdot \text{day}^{-1}$ |
| $d$ | immune cell death rate [59, 60] | $1 \cdot 10^{-2} \text{ day}^{-1}$ |
| $m$ | maximum immune cell proliferation rate | $-10^{-6} \text{ day}^{-1}$ |

To model treatment, two possible procedures were considered. First, an increase in the ability of immune cells to detect and eliminate cancer cells — their *killing rate or efficacy*. Second, adoptive immune cell transfer, corresponding to the injection of immune cells into the system [44, 45]. Both of these mechanisms enhance the suppression of cancer growth by the immune system and can be applied in concert as *combination immunotherapy*. The time at which the killing rate is first enhanced, the treatment time *τ*_*k*_, can differ from the time at which cancer-specific immune cells are first injected into the system *τ*_*E*_. Also, the time period during which each of these treatment approaches are administered can vary, with Δ*τ*_*k*_ the treatment period for killing efficacy enhancement, and Δ*τ*_*E*_ the period for immune cell transfer.

We assume that treatment always consists in the administration of either immunoactivating compounds or immune cells into the host system, and we denote the amount of compound delivered as the administered *dose*. In the increased killing efficacy approach, we assume that the alteration induced by the administration of the compound is permanent, which is reflected in a change of the killing efficacy parameter of NK or CTLs, Δ*c*, or Δ*k*, respectively. The change occurs gradually over the time course of the treatment. For example, an initial CTL killing efficacy *k* before treatment initiation will by increased by Δ*k/*Δ*τ*_*k*_ every day, leading to a final efficacy of *k* + Δ*k*.

In the immune cell transfer approach the change in immune cell numbers is not permanent. Because the immune cells are assumed to be rendered ineffective at a predefined rate (for example by the shedding of NKG2D ligands such as MIC-A, MIC-B [9] or alternatively, by the upregulation the ligands PD-L1 or PD-L2 [15, 16]), immune cell numbers will be affected by already present cancer cells. Thus, immune cell numbers will change depending on the state of the cancer due to its suppressive effect on immune cell proliferation. Conversely, cancer cell numbers will vary due to immune cell killing. Analogously to the dosage of killing efficacy increasing compounds, effector cells are administered at daily doses of Δ*E/*Δ*τ*_*E*_ cells, until the full dose of Δ*E* has been dispensed.

## RESULTS

### Base-model of cancer-immune system interactions

We develop a base model of cancer-immune cell interaction. The model follows a large body of theory that uses two-equation deterministic ODEs to describe the interaction between cancer tumor cells and immune system cells [2, 8, 27–31]. This model aims to replicate some basic features of cancer dynamics with a minimum of added complexity. With such an approach, qualitative insights about the behavior of the system can be obtained by relatively simple mathematical analysis. To this end, we make four fundamental assumptions. First, we assume the existence of immune cells, which are able to detect and kill tumor cells [10, 11]. These cells may eliminate microscopic tumors before they grow to endanger the organism; a process termed *immunosurveillance* [13, 14]. Second, these immune cells comprise the action of all cells that control tumor growth by antigen recognition and subsequent elimination [2], including natural killer (NK) cells [61] and CD8^+^ T cells (or CTLs) [62] and are termed *effector cells*. The process by which the cancer cells are neutralized is called *lysis* [12]. A background level of effector cells is present at all times [63]. Third, we assume that tumor growth is well described by a logistic growth in the absence of immune cells [2]. Fourth, the interactions between tumor and cancer cells are governed by mass-action kinetics [8].

The third assumption of logistic growth warrants special discussion. The dynamics of tumor growth remain a debated issue in the literature. Benzekry et al. compared different theoretical growth dynamics in lung and breast tumor data of mice [64]. They concluded that Gompertzian growth typically best predicts data. Gompertzian growth dynamics are motivated by the observation that tumor growth prior to detection appears to be faster than after detection [2]. This suggests that the initial unbounded growth may be limited by the exhaustion of growth resources or cancer cell’s mutual growth impairment. Gompertzian growth dynamics, developed on the basis of birth and death processes, account for this behavior, and yield a sigmoidal type of growth curve for cancer cell numbers. However, the Gompertz model suffers from serious drawbacks, while other models did almost equally as well as the Gompertzian in predicting data [64]. Because an upper bound of the proliferation rate is imposed by cell division time, the Gompertz model cannot adequately describe the dynamics of very small tumors [2, 29]. Furthermore, theoretical analysis reveals that Gompertzian growth is at odds with the immuno-surveillance hypothesis, because the immune response is unable to eradicate cancers that grow in a Gompertzian fashion [2, 28]. Thus, given these assessments, we chose a growth model that retains a sigmoidal cancer growth curve shape, namely logistic growth. With this, we preserve the notion of an exhaustion of growth resources. We note however, that alternative growth models may also successfully capture tumor growth patterns.

Our base model differs from most models in the literature in that it combines proliferative and suppressive effects of cancer cells on immune cells in one single term that describes its net effect. In this way, we can analyze how the systems behaves depending on the net effect of these opposing forces.

The equations for the base model are:

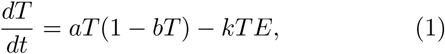

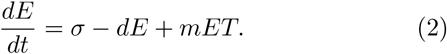

Tumor or cancer cells *T*, grow at a maximal rate *a* in a logistic fashion. The population density of the cancer cells is regulated by the coefficient *b*, which acts as an inverse carrying capacity. The cancer cells are detected and killed by effector cells *E* at a net rate *k* [8]. We assume a constant supply of effector cells at a rate *σ* [34, 35], and a death rate *d* per effector cell [33]. Effector cells can either be stimulated to proliferate or be impaired in their growth at a rate *m*, the net growth increase or decrease due to the presence of cancer cells. In other words, *m* can attain positive as well as negative values.

We proceed to analyze the possible equilibria of this system. We start by observing that 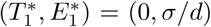 is always a fixed point of the ODEs above. Two further solutions for *T*\* can be represented by the quadratic formula (see *Appendix A*):

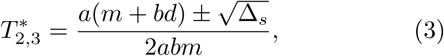

where

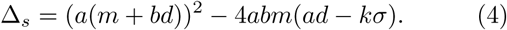

A closer inspection of the properties of the dynamics of (1-2) reveals that all biologically relevant cases, namely those in which 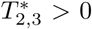, are consistent with *m* < 0 (see *Appendix A*). This corresponds to a net immune cell proliferation suppression by cancer, which can arise by various mechanisms [15, 16, 19–23, 36]. *m* < 0 is also where a bistability pattern in the steady states of (1-2) emerges. The alternative *m* > 0, leads to scenarios which are at odds with well established concepts of cancer modeling, and produce incomplete dynamics (see *Appendix A*). In particular, they conflict with the well-established notion of an *immunological barrier* [8]; the idea that tumors have to grow above a critical threshold to reach a large size close to carrying capacity [2]. Temporary changes in the activity of the immune system can lead to fluctuations in tumor size which place its size above the barrier, which then lead to cancer.

There are three possible cases of sign arrangements of the roots of (3) under *m* < 0: i) both negative, ii) one positive and one negative, iii) both positive. Scenario i) has 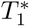 as only biologically plausible solution. Only the cancer-free state exists. Case ii) signifies a single attractive equilibrium at a non-zero tumor size. The emergence of a cancerous cell suffices to ignite a replicative process that induces the establishment of a tumor close to carrying capacity 1*/b*. Thus iii), where 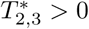, is the only case which admits stable equilibria compatible with an existing immunological barrier.

For 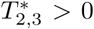 to be satisfied, and while assuming that *a* > 0 and *b* > 0 for biological reasons, we obtain the following conditions (see *Appendix A*):

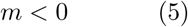

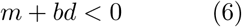

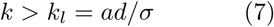

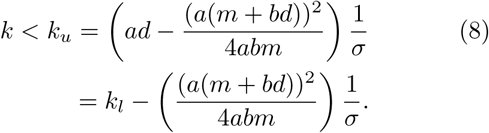

These results reveal a bistability pattern that is mediated by the killing efficacy *k*. Figure 2 shows how the increase in the parameter *k* leads to a bifurcation in the stable states of *T* and to bistability for *k*. At *k* < *k*_*l*_ the system will reside in the aforementioned case ii). By increasing *k* above *k*_*l*_ but below *k*_*u*_, the system will enter case iii), and move to i) as *k* > *k*_*u*_. In line with our expectations, values of *k* below the threshold value *k*_*l*_ represent a similar situation as would be expected in the absence of immune cells: unchecked cancer growth. If *k* is gradually increased above *k*_*l*_, the tumor cells would have to begin replicating at increasingly large initial sizes in order to avoid being absorbed by the attractor at 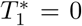, that is, to be suppressed by the immune system. This is where the bifurcation appears, and now three equilibria, of which two are stable, dominate the dynamics. The parameter range spanned between *k*_*u*_ and 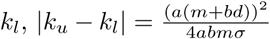 is highly dependent on the ratio of *b* and *m*, as well as *σ*. Lastly, large values of *k* above *k*_*u*_ entail that even few effector cells are able to clear the tumor, and even large tumors are eliminated with certainty.

**FIG. 2.**
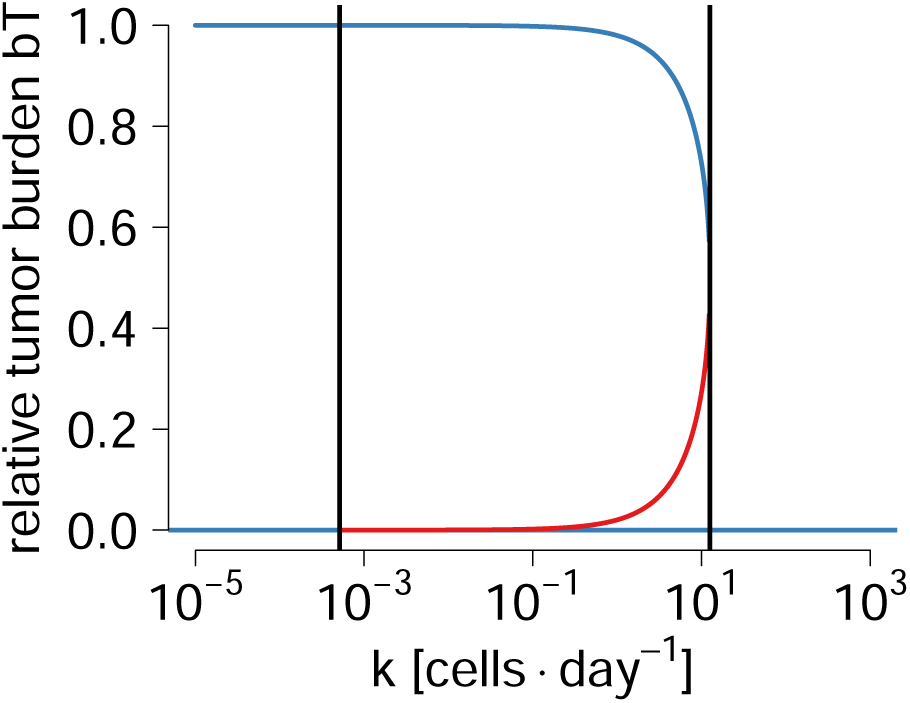
Bistability of immune control of tumor growth. Increasing values of *k* lead to the emergence of a bifurcation in equilibrium tumor cell numbers. The emergence of the bifurcations is indicated by black, vertical dashed lines. The upper blue line shows the largest stable steady states and the lower dashed blue line shows the smallest of the steady states, while red lines show unstable steady states for a given *k* value. If *bT* is one, the cancer has reached carrying capacity. At *bT* = 0, no cancer cells exist. Thus, above a threshold value *k*_*u*_, the cancer is cleared. The parameter values are *m* = −10^−6^, *σ* = 10 [35, 65], *d* = 2 × 10^−2^ [59, 60], *a* = 0.514 [5], *b* = 1.02 × 10^−9^ [5].

At intermediate *k* (case iii) the base level of effector cells crucially affects the dynamics. In the absence of cancer, the equilibrium level of effector cells is at *σ/d*. The appearance of cancer cells will induce a killing process mediated by *k*. High enough effector cell levels will suppress tumor growth. In reality, the tumor might fluctuate above some threshold within which effector cells control it, and the suppression of further T cell proliferation (*m* < 0) by the cancer will dominate over the killing. Thus, the conditions (5-8) mark the region of killing efficacy values that constitute an immunological barrier to the cancer. Increasing *k* values imply that this barrier is heightened: existing effector cells improve their capacity to completely eliminate the cancer.

This analysis may serve as a model to think about how to efficiently combine immunotherapy approaches. These results suggest that one mechanism to generate biologically plausible bistability is consistent with situations in which the cancer’s immunosuppressive effects outweigh its immunoproliferative effects (*m* < 0). The existence of a bifurcation implies that increasing killing efficacies will not simply gradually diminish cancer cell numbers. Instead, an increase above above a critical value *k*_*u*_ can terminally clear the cancer. Equally, if *k* is located in the range *k*_*l*_ < *k* < *k*_*u*_, the adoptive transfer of effector cells may work by perturbing cancer cell numbers below the unstable equilibrium point 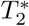, after which it will be absorbed into 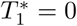 and be cleared.

### Model variations

The here presented base-model, despite being strongly simplified, can help us intuitively understand more complex models of cancer-immune system interactions. In the following, we demonstrate that the basic bistability phenomenon can be replicated in two related, but more complex variations of the base-model. When possible, we analyze whether multistability can arise, because of its relation to *cancer dormancy*, and because it might mitigate the effect of treatment.

### Incorporating saturation effect of tumor cells on immune response

A natural way to extend the base-model is to include more biologically plausible assumptions about the behavior of effector cells. Here, we retain the basic assumptions going into the base model, but refine the way that effector cells are reacting to the presence of tumor. In particular, we follow the approach by Conway et al., which allows for a very general set of behaviors of effector cells to arise and also provides estimates for the parameters [36]. This study is set in the context of HIV, where a great body of theory has been devoted to the particulars of CTL behavior [33]. The equations are:

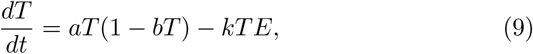

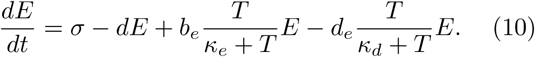

Here, the interactions governing the rate of change in tumor cells have remained intact. Effector cell growth can now be stimulated at a maximum rate *b*_*e*_ [36, 66]. The proliferation rate saturates with the number of tumor cells *T*, and is half-maximal at the Michaelis-Menten constant *κ*_*e*_ [36]. In contrast, effector cells can be exhausted by contact with tumor cells, and die from its consequences at a maximal rate *d*_*e*_ [36, 67]. As with proliferation, a saturation ensues in the exhaustion which is mediated by *κ*_*d*_. For typical values of these parameters see Table II in *Appendix B*. Models with a saturation term 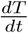 have been analyzed before [68], and have been thoroughly discussed in for example [3].

**TABLE II.**
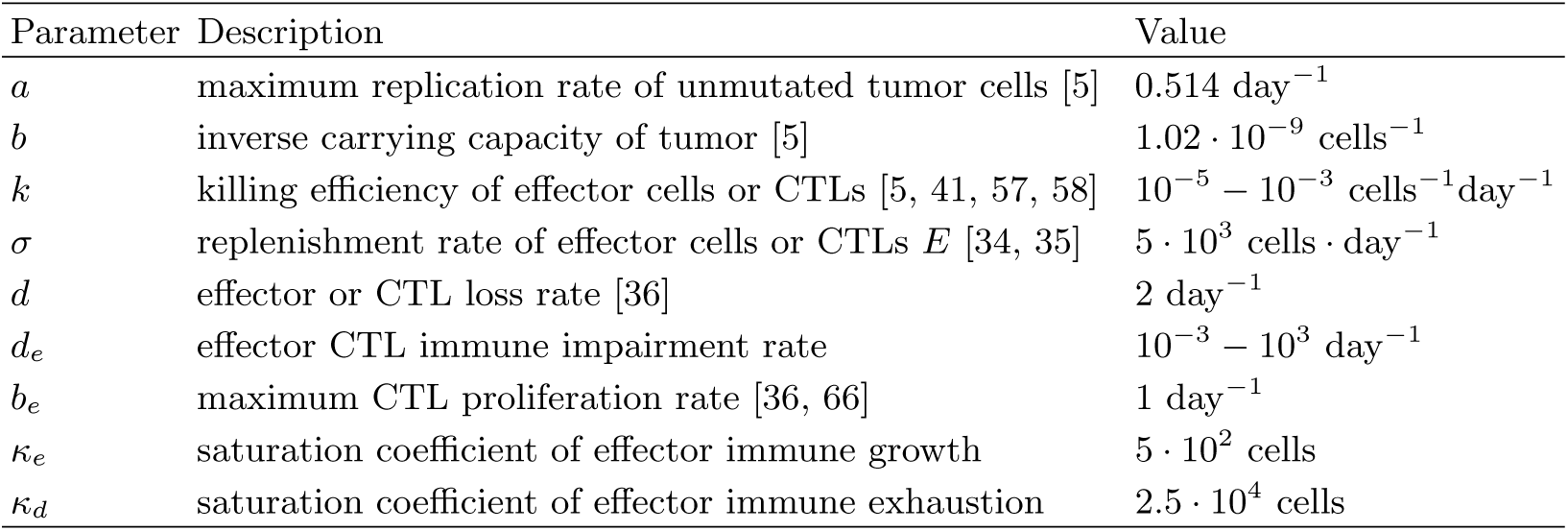
Parameter values for the extended base model with saturation (9-10).

| Parameter | Description | Value |
| --- | --- | --- |
| $a$ | maximum replication rate of unmutated tumor cells [5] | $0.514 \text{ day}^{-1}$ |
| $b$ | inverse carrying capacity of tumor [5] | $1.02 \cdot 10^{-9} \text{ cells}^{-1}$ |
| $k$ | killing efficiency of effector cells or CTLs [5, 41, 57, 58] | $10^{-5} - 10^{-3} \text{ cells}^{-1} \text{ day}^{-1}$ |
| $\sigma$ | replenishment rate of effector cells or CTLs $E$ [34, 35] | $5 \cdot 10^3 \text{ cells} \cdot \text{day}^{-1}$ |
| $d$ | effector or CTL loss rate [36] | $2 \text{ day}^{-1}$ |
| $d_e$ | effector CTL immune impairment rate | $10^{-3} - 10^3 \text{ day}^{-1}$ |
| $b_e$ | maximum CTL proliferation rate [36, 66] | $1 \text{ day}^{-1}$ |
| $\kappa_e$ | saturation coefficient of effector immune growth | $5 \cdot 10^2 \text{ cells}$ |
| $\kappa_d$ | saturation coefficient of effector immune exhaustion | $2.5 \cdot 10^4 \text{ cells}$ |

As with the base model, 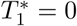 is always a fixed point. The rest of the fixed points are determined by the roots of a cubic equation (see *Appendix B*). For the system of equations to generate bistability, that is, to give rise to exactly two positive equilibria in *T*, the following set of conditions needs to be satisfied (see *Appendix B*):

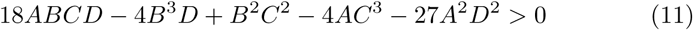

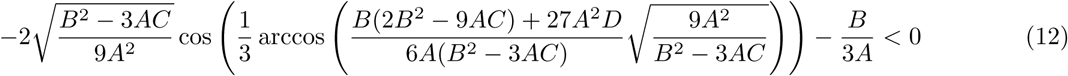

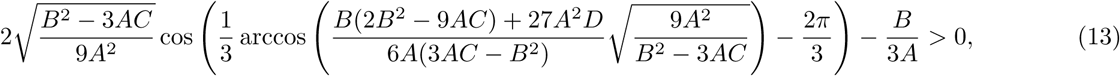

where

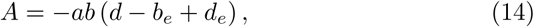

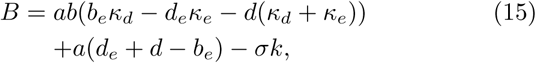

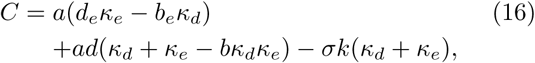

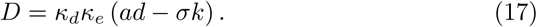

Analogously to the base model, a closer inspection of this result reveals that biologically plausible dynamics are consistent with *A* = *ab* (*b*_*e*_ − *d*_*e*_ − *d*) < 0 (see *Appendix B1*). Interestingly, this expression is independent of *κ*_*e*_ or *κ*_*d*_. Since *a, b* > 0, this implies that *b*_*e*_ < *d*+*d*_*e*_0 which is analogous to the situation where *m* < 0 in the base model. The treatment rationale identified in the base model may therefore also be applicable for the extended base model with saturation (9-10).

In Conway et al’s work [36], this condition is satisfied by *b*_*e*_ < *d*_*e*_, which is also functionally equivalent to *m* < 0 in our base-model: the effector cell population decreases due to cancer-mediated exhaustion. Importantly, the parameter choice in Conway et al. is also consistent with the emergence of bistable and multistable equilibira.

Besides bistable patterns, the system (9-10) can also generate a pattern of multistability. The conditions to obtain four equilibria in *T* for (9-10) reads:

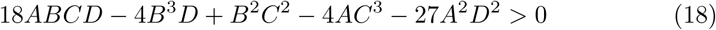

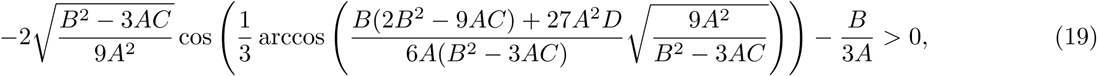

where *A, B, C* and *D* are as in (14). Again, a biologically plausible arrangement of equilibrium points is in agreement with *A* < 0. Figure 5 shows that bistability becomes common for *d*_*e*_ > 1 and *k* > 5 · 10^−4^, with the elimination of cancer ensuing after a critical *k*-threshold is surpassed.

The existence of multiple stable equilibria may be interpreted as *cancer dormancy* (see *Discussion*). Cancer dormancy is the phenomenon of a period of non-growth of tumors. Often, this occurs in small, nearly undetectable tumors residing within body tissues [69]. These small tumors are said to be dormant, that is, not growing to large and more threatening sizes. Figure 5 shows that multistability is possible in (9-10). The existence of multistability in two-equation models has been predicted and shown in other work [8, 29, 70]. Here, we give precise analytical conditions for its emergence under (9-10). When in a multistable regime, an increase in killing efficacy *k* might not directly lead to the elimination of the cancer if treatment is started when the cancer is near carrying capacity 1*/b*. Instead, a new microscopic steady state (MISS, [29]) might be attained before a further increase in *k* leads to cancer clearance.

### Incorporating natural killer (NK) cells and tumor-specific CTL response

In a next step, we incorporated a further level of complexity by distinguishing between two types of effector cells: natural killer or NK cells, and cytotoxic T lymphocytes or CTLs. Models that account of the different roles between NK and CTL can be highly complex [5]. To better understand where possible bistabilities originate from, we restrict ourselves to an extension of the base model with saturation, following an approach inspired by de Pillis et al. [5] and Conway et al. [36]. Effector cells are now split into NK cells *N* and CTLs *E*:

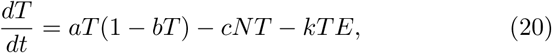

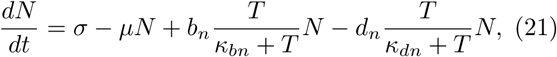

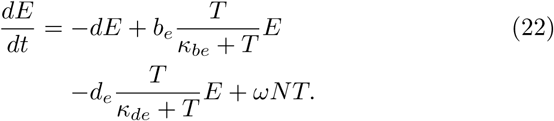

Here, cancer cells *T* are killed at rates *c* by NK cells, and at rates *k* by CTLs. The dynamics of the NK is now analogous to the dynamics of immune cells in (9-10). We assume a constant supply of NK cells *σ* stemming from the host’s hematopoesis. NK cells die naturally at a rate *µ*. The maximum NK proliferation rate induced by the presence of cancer cells is *b*_*n*_, and the saturation coefficient is *κ*_*bn*_. Again, exhaustion occurs at a maximum rate *d*_*n*_ and the saturation in *T* is half maximal at *κ*_*dn*_. For CTLs, following [5], we assume that there is no constant supply of cells. Instead, CTL growth is stimulated by the NK-cancer cell interactions at a rate *ω*. This modeling approach ensures that CTLs are activated only after the emergence of the NK immune response [5]. Several biological mechanisms appear to exist by which NKs can stimulate CTL growth [71]. The work of Fan et al suggests that already activated NK cells can facilitate the priming of CTLs by means of IFN-*γ* [72]. The proliferation and exhaustion terms are as in the extended base model.

The model (20-22) can generate substantially more complex behavior than the two previously analyzed. As in the previous models (1-2 and 9-10), 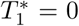 is always a fixed point. However, the NK-CTL model may allow for up to five other fixed points (see *Appendix C*). This is because finding the steady states of (20-22) can be reduced to finding the roots of the rate of change of *T*, 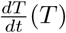, which is a polynomial of sixth order. The five fixed points besides 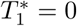, are the roots of a fifth order polynomial, for which no general solutions exist. Thus, we cannot draw similar conclusions for the existence of real fixed points as in the previous, relying on analogous conditions on discriminants Δ_*s*_ > 0 or Δ > 0. However, similarly to the base model (1-2), a similar condition to *m* < 0 is compatible with biologically plausible arrangements of fixed point’s stabilities. The expression analogous to *m* < 0 is (see *Appendix C*):

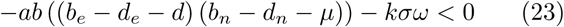

The analogy to *m* < 0 arises from *k, σ, ω* > 0, valid bounds in most biological contexts. If tumor cells are able to exhaust NKs and CTLs, that is, if both (*b*_*e*_ − *d*_*e*_ − *d*) < 0 and (*b*_*n*_ − *d*_*n*_ − *µ*) < 0, then the system can display biologically reasonable behavior. Note that unlike the previous models, this condition can be satisfied by other means as well, such as increasing *k*. Again, the condition is independent of the four saturation coefficients *κ*_*be*_, *κ*_*de*_, *κ*_*bn*_, *κ*_*dn*_.

Due to the analytical unfeasability of the model (20-22), we resorted to numerical methods to prove the existence of basic bistability patterns (see *Appendix C*) [50, 51]. We found that the system is able to display bistability similar to that found in the extended base model with saturation (see Figure 6).

### Bistability-based Strategies of Cancer Immunotherapy

The existence of bistability patterns in simple non-spatial cancer models as well as its variations, can be informative to the assessment of immunotherapeutic options and of their efficacy. Taking the base model (1-2) as a foundation, three intervention approaches seem apparent. First, the elimination of the exhaustive effects of cancer on the immune cells (*m* < 0 → *m* > 0). Second, the increase of the killing efficacy of effector cells above some threshold (*k* < *k*_*u*_ → *k* > *k*_*u*_). Third, the administration of effector cells (*E* → *E* + Δ*E*).In terms of the dynamics, this represents pushing the state of the system into the attraction basin of *T* = 0. Combinations of these therapy approaches have previously been explored in simulations [68], and constitute a promising avenue of future research [9].

In current immunotherapy, the two main available tools for cancer cell reduction correspond to the second (*antibody therapy*) and third (*adoptive T cell transfer*) options [9, 26]. In antibody therapy, an increase in killing efficacy is attained by disrupting cancer cells mechanisms to avoid recognition by immune cells, effectively impairing T cell killing. This impairment occurs by the acquisition of mutations in cancer cells that, for example, lead to the expression of the PD-L1 and PD-L2 ligands [15, 16]. These ligands are known bind to the PD-1 receptors on T cell surfaces, thereby downregulating the activation of the T cells. Monoclonal antibodies binding to the ligands can interrupt this cancer escape mechanism and make the cancer cells visible for the immune system again, making them a useful way to effectively increase *k*. In *adoptive T cell transfer* [44, 45], T cells are pre-programmed to kill host cells that carry particular biochemical signatures, for example certain peptides on their surface. The signatures are chosen such that they match characteristic features of cancer cells. Subsequently, these specific T cells are grown and injected into the blood stream of the patient [9].

These two novel treatment methods can be combined to take advantage of the bistability phenomenon in cancer. We used the base model (1-2) to investigate how a combination of both approaches could be used to clear the tumor, while accounting for the stochastic effects arising from singular cell-to-cell interactions. Starting from an already established tumor, increasing the efficacy of killing of effector cells will not by itself necessarily lead to the elimination of the tumor, unless very high levels of killing efficacy can be attained. Instead, increasing the killing efficacy by two orders of magnitudes will lead the system to equilibrate at tumor cell numbers lower than the carrying capacity (see Fig. 2). If the treatment with monoclonal antibodies has been effective enough, it will have shifted the system into a regime with two stable equilibria, out of which one is the cancer-free state. If now the system is perturbed further with adoptive T cell transfer into the attraction basin of the cancer-free equilibrium, the waning of the effects of the first treatment will not lead to the reemergence of the cancer. Thus, the generation of a temporary bistability in the system can be exploited to perturb it into a cancer-free state.

To model the combined effects of killing efficacy increases by PD-1 specific monoclonal antibody and adoptive T cell transfer treatments, we used stochastic version of the base cancer immune interaction model (1-2) (see *Materials and Methods*). Figure 3 shows the time courses of the dynamics with and without treatment. Without treatment, a cancer that has surpassed the immunological barrier will grow unrestricted up to levels very close to carrying capacity (Fig. 3A). The combined administration of effector cells and killing efficacy-increasing compounds at first only gradually reduces cancer numbers (Fig. 3B). Daily administered effector cells Δ*E/*Δ*τ*_*E*_ (*Materials and Methods*) can only temporarily affect the dynamics before they are rapidly suppressed and exhausted by the cancer (*m* < 0). When the state of the system is pushed into the attraction basin of the cancer-free state, cancer numbers rapidly plummet towards zero. Further injections of effector cells become unnecessary, and effectors build up.

**FIG. 3.**
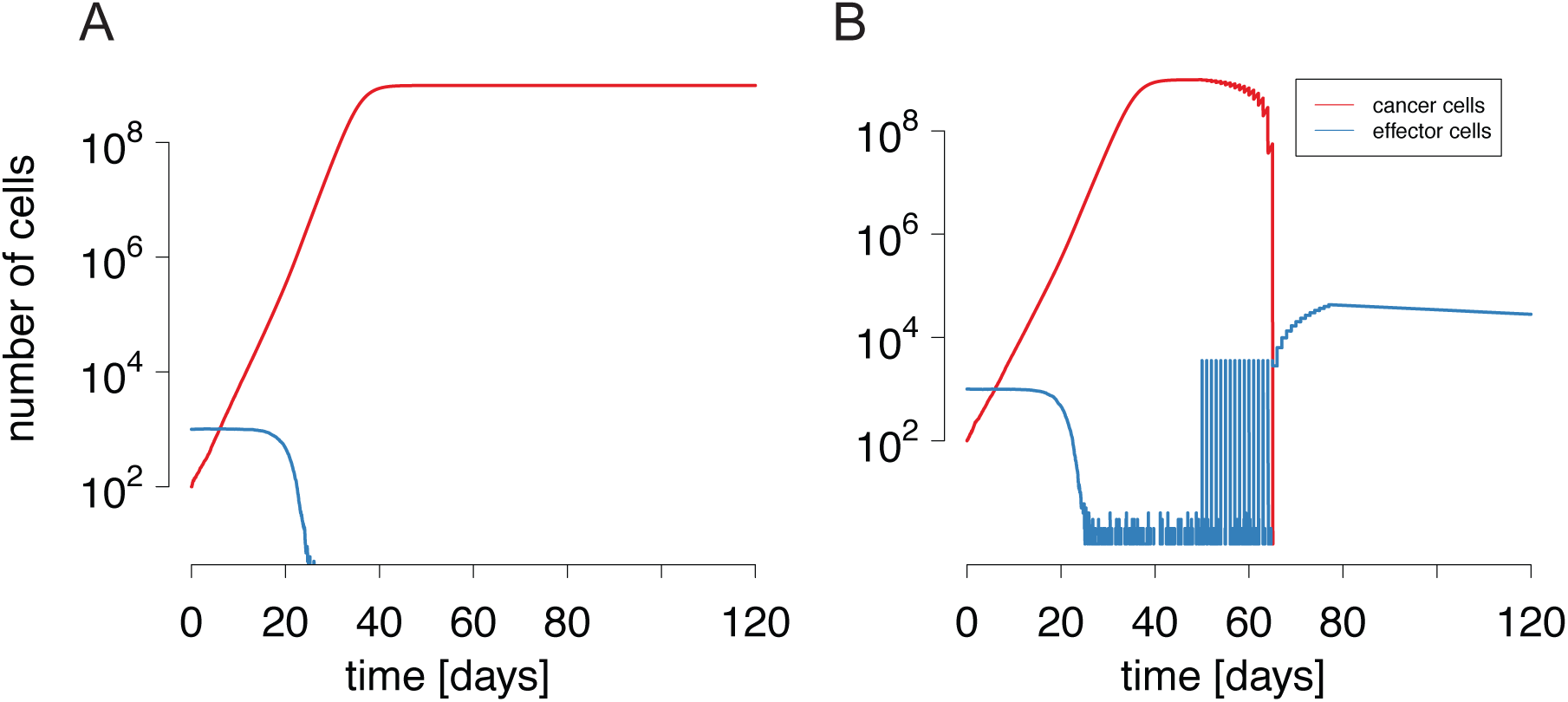
Simulation runs of the base model without A) and with B) immunotherapeutic treatment. A) Cancer replication begins at *T*(0) = 10^2^, and the natural equilibrium of the effectors is at *E*(0) = 10^3^. The cancer grows to carrying capacity in a time frame of around 45 days. B) Combined immunotherapeutic treatment is initiated at 50 days after the the cancer has begun to grow. It lasts for 21 days in antibody therapy, and 28 days in adoptive cell transfer. Killing efficacies are increased to Δ*k* = 10^−1^, while a total of Δ*E* = 10^5^ cells are injected in a gradual fashion. Immune cells are eliminated rapidly by the powerful immune exhaustion effects exerted by the cancer cells. Parameters values are as in Table I. In particular, *k* = 10^−4^.

We then investigated whether an increase in *k* and the administration of effector cells *E* work together in a synergistic or antagonistic fashion to remove the tumor. Figure 4 shows the outcome of combination immunotherapy, initiated simultaneously for antibody and T cell injections. Each immunotherapeutic approach may clear the cancer on its own. A marked frontier between cancer presence and clearance emerges. A lowering of killing efficacies along this frontier will lead to insufficient pressure to clear the cancer, but can be compensated by an increase in adoptive transfer doses. The linear shape of the frontier indicates that the two approaches do not mutually impair each others’ function.

**FIG. 4.**
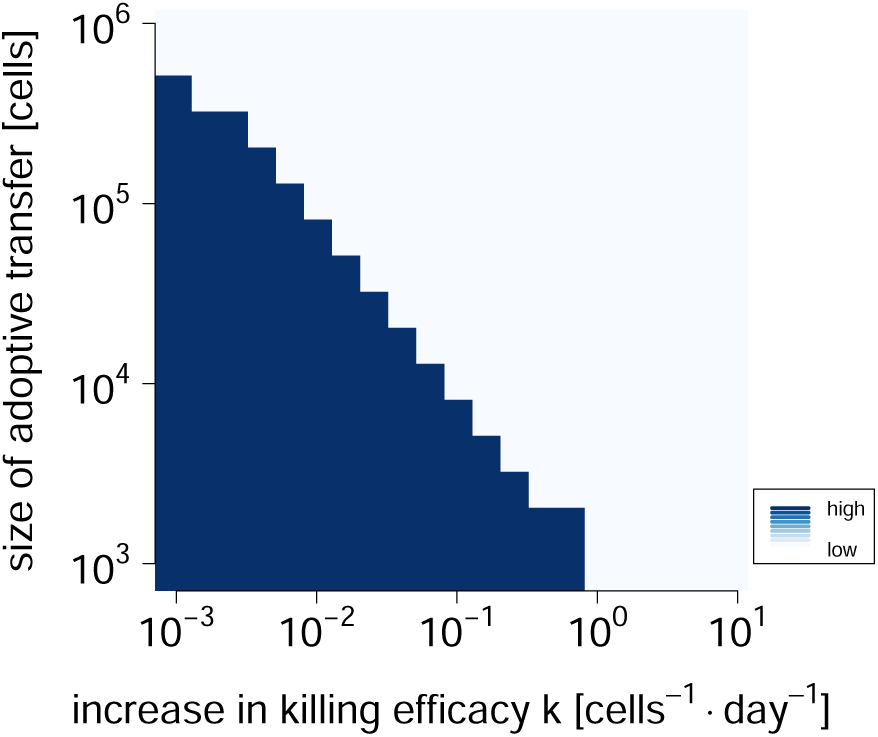
Cancer cell numbers after combination immunotherapy after 120 days. Immunotherapy begins at day 50, after the cancer has grown to full size, and lasts for a single day. Combination immunotherapy is implemented by increasing the killing efficacy of the effector cells and adoptive immune transfer of effector cells. Darker colors indicate high cancer cell numbers, implying that the cancer persists, while lighter colors indicate low cancer cell numbers. Parameter values are as in Table I.

## DISCUSSION

In this study, we have shown that the base model can only reproduce biologically plausible behavior if the suppressive effects exerted by cancer cells on immune cells dominate their proliferative effects. Under these circumstances, the base model displays a conspicuous pattern of *bistability*: The cancer-immune interaction dynamics gives rise to two distinct, stable states (a cancer-free, and a full-grown tumor state). Under bistability, the modification of the killing efficacy can lead to a bifurcation in cancer cell numbers, where the system may abruptly be tipped into a new, cancer-free state. Furthermore, in situations where exhaustion prevails over proliferation in immune cells, all analyzed models can produce bistability patterns that are biologically plausible. If this condition is not satisfied, the base model cannot produce biologically plausible behavior across a range of *k*. We also formulated more complex extensions of the base model, which can generate *multistability*, a dynamic behavior that can be interpreted as stable microscopic cancers [28], or *cancer dormancy* [8, 69]. We gave the exact conditions under which multistability might arise in one model extension inspired by [36]. We also examined how bistability may be used for effective combination immunotherapy. We tested a combination of two different immunotherapeutic approaches in stochastic simulations of the base model. We found that the combination of treatment interventions is able to clear the cancer, and that the different treatments approaches do not impair one another.

How both treatment approaches investigated here would work in isolation has to some extent been studied in other work [2, 27, 29]. However, it is less apparent that they may be employed to work in concert without mutual impairment. This is important when considering side effects of these therapies. For example, the administration of PD-1 antibodies in mice have resulted in lung inflammation and cardiomyopathy [9, 73, 74]. Thus, we find that combination immunotherapy may help minimize the risks associated with standalone approaches.

Another advantage of this study lies in that it specifically shows how multistability may arise from standard assumptions about cancer-immune system interactions (saturation terms), and in that it gives precise conditions for its emergence. Although the first extended model with saturation (9-10) is not intended to admit multiple steady states, these arise as a consequence of the basic assumptions about interactions.

Intermediate-sized cancers in multi-stable regimes may be interpreted as cancer dormancy [8, 29, 70], but the model ((9)-(10)) does not explicitly explain how they may escape immune control. For immune escape to arise, additional processes must be assumed, such as stochastic perturbations or immunoediting [75]. A discussion of how tumors might rise to large numbers in models very similar to the base model with saturation has been given in Kuznetsov et al. [8] and by Wilkie and Hahnfeldt [76]. In the Kuznetsov et al. model, a separatrix between the two main attraction basins passes close by the trivial, cancer-free steady state. When a cancer arises and starts to replicate at low numbers, it should follow a trajectory into a dormant, stable steady state. However, stochastic fluctuations can push the system’s state into the other attraction basin.In the Wilkie and Hahnfeldt model, the saddle point is the dormant state itself. Another of the main mechanisms hypothesized to drive the transition from dormancy to large tumors is *immunoediting* [69]: The prolonged growth suppression of the tumor by the immune response leads to the selection of cancer mutations that escape immune pressure, effectively reducing the immune killing efficacy *k* [75]. The model (9-10) can also offer an intuitive explanation for this process, whereby a smaller, undetectable and stable equilibrium of cancer cells is maintained by a relatively weak immune response. The further decrease of the immune response efficacy by means of immune escape processes leads to the establishment of a full grown tumor. This is exemplified by the fact that decreasing *k* in Figure **??** pushes the system into a region of parameter space where there exists a stable steady state for the tumor at carrying capacity, that is the full grown cancer state.

Most other models of cancer-immune interaction so far have attributed the phenomenon of dormant states to the existence of an additional compartment: quiescent cancer cells [69, 77]. These cells are assumed to replicate at a slower rate than normal cancer cells, and can revert back to a fast growing state by means of phenotypic switching or by acquiring further mutations [69]. In line with other work, the model (9-10) explains the existence of dormancy by a specific balance of cancer cell growth and killing attained in cancer-immune interactions, without relying on any additional compartments or assumptions. A notable example of how dormancy can emerge from cancer-immune interactions alone is given in Kuznetsov et al. [8]. A third mechanism for the emergence of dormancy has been given by [76]. In this mechanism, dormant states are represented by saddle nodes traversed by a separatrix demarcating the adjacent attractor regions of either growth progression or tumor clearance.

This point emphasizes a last advantage of non-spatial ODE models: Understanding cancer growth requires an appropriate description of cancer-immune system interactions at multiple scales [1]. These scales range from cancer microenvironments to large numbers of already systemic cancers. ODE models offer useful tools to combine the behavior of both into a single framework [2], accounting for the frequency-dependent growth at early stages as well as the dominant immunosuppressive effects achieved by cancer when approaching carrying capacity levels.

A major caveat of the base model is that it cannot elicit an immune response. The elicitation of an immune response by a cancer, with a subsequent rise in effector cells is an important aspect of cancer-immune system interactions. To achieve this, the base model would have to include a proliferation term for effectors that takes on different values than the suppression in the *T, E*-plane. The base model is thus more useful to study the aspects of how immunotherapy can be deployed to return the CISI system below an immunological barrier by external perturbation.

While simple modeling frameworks offer greater possibilities for an in-depth understanding, this approach also has its limitations. For instance, we have largely neglected stochastic attributes of cancer-immune system interactions in our mathematical analysis. These may mostly arise from the discreteness of cell-to-cell interactions, and are well captured by simulating the models by a Gillespie algorithm [54]. In this study, we have addressed this shortcoming by adopting a stochastic simulation framework to implement immunotherapy. Other forms of stochasticity —for example the random accrual of malignant mutations in cancer cells— are also not explicitly modeled. Instead, they are assumed to be captured by model parameter values. Environmentally based fluctuations [2, 78], or changes in the exerted immune pressure due to, for example, disease [79], are also neglected.

The models here analyzed do also not account for spatial structure (discussed in more detail in [80–82]). Spatial structure may change the way that effector cell killing affects cancer growth, as well as how the presence of cancer cells may mediate immune cell proliferation. In particular, we did not explore the fractional cell kill laws as introduced by de Pillis et al. [5], hypothesized to account for some of the geometrical features of tumors [31]. In this approach, the total killing exerted by effectors *K*(*E, T*) is governed by the dePillis-Radunskaya-Wiseman (PRW) law [31], where *K*(*E, T*) = *D*(*E, T*)*T* and 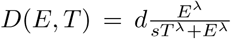. Structurally, how [5] implement the recruitment of NK cells differs only slightly from our implementation in (20). However, how CTLs are recruited differs in structure from the simpler terms analyzed here in our models. With λ < 1 (obtained from model fits to mouse data [31]), the behavior of the recruiting is qualitatively similar to that studied here: the recruiting of CTLs would then saturate with increasing cancer cell numbers *T*, but continue growing with increasing effector cell numbers *E*.

We thus assume that while the PRW law introduces an advantageous new concept in the modeling in tumor-immune system interactions, our deviating from it will not yield marked qualitative differences. Instead of attempting to capture tumor geometry behavior, the models analyzed here are rooted in the tradition of virus dynamics —especially HIV— which assumes well-mixed cell types [32, 36]. Thus, the PRW law seems to mainly address problems arising from tumor geometry, while this study focuses mainly on systemic cancer types —cancer types that do not manifest in single tumors only [2].

We have also not included the action of cytokines in our analysis, which are typically accounted for with a separate, additional equation [2, 68, 83]. We have therefore not been able to assess the effectiveness of cytokine-based immunotherapy approaches in combination with the ones studied here. Models with cytokines display features like the persistence of large tumors, tumor dormancy, and tumor clearance upon immunotherapeutic treatment, as well as oscillations between these states [2, 68]. Including cytokines into a more comprehensive modeling framework would be an interesting topic for future work.

Our study has to be interpreted in the context of other models of cancer-immune system interactions. The most comprehensive mathematical analysis of two-equation models has been put forth by d’Onofrio [28, 29]. d’Onofrio analyzed a generalized mathematical model in two variables —*x* denotes cancer and *y* effector cell densities—, deriving some general results on the existence of steady states and cancer eradication given some general mathematical conditions on the interactions between these two cell types. Solutions were provided for the generalized model, but except for the rate of adoptive transfers *θ*(*t*) (where *t* is time), no specific dependence was given on how steady states change with parameter value modifications. Our base model corresponds to a special case of his general model (equations (1-2) in [29]), with *ϕ*(*x*) = *k, f* (*x*) = *a*(1 − *bx*), *β*(*x*) = 0, *q*(*x*) = 1 and *µ*(*x*) = (*d* + |*m*| · *x*). As in the models investigated in this study, d’Onofrio has observed that his generalized model admits a cancer-free state, and also predicted that it may attain multiple stable equilibria, which he interpreted as microscopic steady states (MISS) and which we interpret in the context of cancer dormancy. Our own results thus confirm some of d’Onofrios, but go further to explore how specific cancer-immune system interaction models are concretely affected by changing dynamical properties that may be tailored for immunotherapy. In particular, we wanted to explore some of the properties of CISI models that underpin the mechanisms that may give rise to bistability patterns (the aforementioned dominance of immunosuppressive effects). In our models, we were therefore more interested in explicit analytical results, which would allow us to study how bistability patterns depend on effector cell killing efficacy. We also extended this approach to include how adoptive transfer might function under conditions with stochasticity.

A similarly comprehensive analysis of how CISI models may give rise to successful adoptive immunotherapy treatments has recently been put forth by Talkington and colleagues [3]. Similarly to our own conclusions, Talkington et al. identify bistability as a major prerequisite for successful adoptive immunotherapy. Their approach is also to review a series of models of increasing complexity, whereby complexity is understood to represent incorporations of additional aspects of the immune system, such as helper cells, interleukin and naïve T cells into a base model. The base model is Kuznetsov et al.’s early model from 1994 [8], which, akin to our base model, assumes only two compartments: tumor and effector cells. The Kuznetsov model allows for bistability, with a stable state of the tumor close to cancer eradication. For all other models, Talkington et al. show that when they can give rise to bistability, adoptive immunotherapy leads to successful outcomes in simulations. Future work could address whether the mechanism for bistability emergence identified in this study, the dominance of immunosuppression by cancer over immune cell proliferation, is also the one that gives rise to bistability in the models examined by Talkington et al.

Future work could address whether the mechanism for bistability emergence identified in this study, the dominance of immunosuppression by cancer over immune cell proliferation, is also the one that gives rise to bistability in the models examined by Talkington et al. To this end, it will be useful to embark on a more comprehensive analysis of how mathematical models as the ones put forth here, bring about some behavior of interest, such as bistability. One approach that could be taken in this direction follows the axiomatic modeling framework pioneered by Komarova et al. [84] and d’Onofrio [28, 29].

In a departure from the work of de Pillis et al. [5], we have concentrated on model features typically used in HIV modeling, borrowing in particular from the study of Conway and Perelson [36]. The reason for this choice is that ODE-based modeling has a long tradition in HIV and virus dynamics modeling [32, 33, 85]. A great wealth of data have helped to validate interaction terms of different cell types in mixtures, and particularly, how CTLs kill. In our view, these advantages can be fruitfully employed in cancer modeling as well. A more inter-disciplinary integration, in particular with respect to CTL behavior, will benefit both fields, and allow for the analysis of structural similarities between models that might be harnessed for immunotherapy design.

Our models show that biologically plausible cancer-immune system interactions may be utilized to induce cancer-free states. Increases in the killing efficacy of immune effector cells can destroy the bistability pattern inherent in those models, abruptly removing the basis for cancer growth.

## ACKNOWLEDGMENTS

We are indebted to Gabriel Leventhal and Elias August for useful discussions.

## APPENDIX A: MATHEMATICAL ANALYSIS OF THE BASE MODEL

Here, we describe the analysis of the system of equations (1 - 2) and its equilibria (termed *E*\* and *T*\* in the following). To this end, we first solve *dE/dt* = 0 for *E*\*, and obtain an expression in *T*\*. We then subsitute *E*\* in 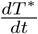, and obtain a polynomial in *T*\*. We aim to find the roots of 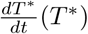, which are the fixed points of the system of equations. Removing the one obvious fixed point given by 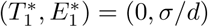, we are left with a quadratic equation in *T*\*. To obtain the other fixed points, we analyze its roots:

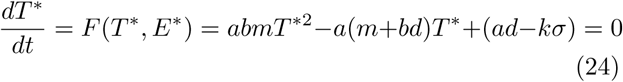

The solutions of (24) are given in (3).

With this, we can derive the exact conditions for (i) both 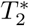 and 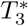 below zero (ii) 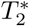 and 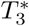 of different signs (iii) both 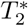 and 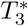 above zero.

We go through all cases, starting with case i). For both solutions 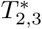 to be negative, we have two subcases ia) and ib) of sets of conditions. Either (ia), we require that 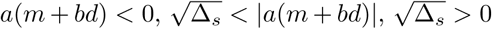 and *abm* > 0, or equivalently (ib), 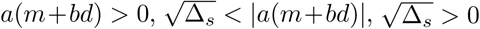 and *abm* < 0.

We want to investigate the similarities sub-casesto simplify the expressions that lead to i). Let us start by analyzing the condition 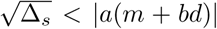, which is the same in both sub-cases. This is equivalent to stating that 4*abm*(*ad* − *kσ*) > 0. In this expression, *a, b, d* and *σ* have to be assumed to be positive for biological reasons. *k*, the net effector cell killing of cancer cells can theoretically become negative, which could be the consequence of cancer growth stimulation by the presence of effector cells as reported in some very recent studies [9]. The net growth stimulation from cancer cells *m* could theoretically become negative also, since cancer cells are known to evolve mechanisms that activate suppressive T regulatory cells and thus escape effector cell action [9]. This immunosuppressive effect could overpower the growth stimulation of T cells induced by the presence of cancer. Thus, for 4*abm*(*ad* − *kσ*) > 0 to be valid, there exist two options: Either *ad* − *kσ* > 0 and *m* > 0, or *ad* − *kσ* < 0 and *m* < 0.

The other condition that appears in both sub-cases is Δ_*s*_ > 0, which is required to obtain real solutions for 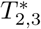. Depending on the sign of *m*, this translates into different conditions for *k*: if *m* > 0, then *k* > *ad* − (*a*(*m* + *bd*))^2^*/*4*abm*)/*σ*. and otherwise if *m* < 0, *k* < (*ad* − (*m* + *bd*))^2^*/*4*abm*)/*σ.*

These conditions on *m* must concord with the conditions on *abm* derived before, since *a, b* > 0. And thus if *m* > 0, we are in the same sub-case as *a*(*m* + *bd*) < 0 and *abm* > 0, whereas *m* < 0 must coincide with *a*(*m* + *bd*) > 0 and *abm* < 0.

Thus, to obtain 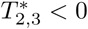, we have two sets of conditions dependent on parameter *m*. The first set of conditions is:

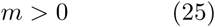

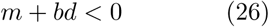

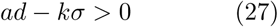

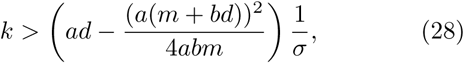

whereas the second set of conditions reads:

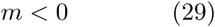

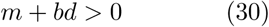

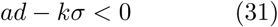

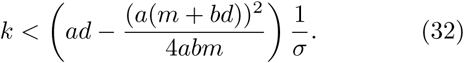

Case ii) is of greater interest, since it entails that there will be one other positive solution to (1-2). As in case i), there exist two situations in which 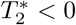 and 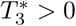 is attained. For these conditions to be both true, it follows that either 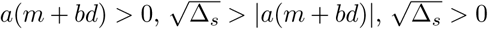 and *abm* > 0, or equivalently, *a*(*m* + *bd*) < 0, 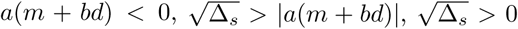 and *abm* < 0. Note that compared to i), the inequality that compares |*a*(*m* + *bd*)| with 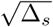 has been inverted. Fortunately, again, the inequalities on |*a*(*m* + *bd*)| are equivalent in both sub-cases, so we can proceed analyzing it.

Following an analogous discussion as in i), it follows that for 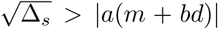 to be true requires either *m* < 0 and *ad* − *kσ* > 0 or conversely, *m* > 0 and *ad* − *kσ* < 0. The conditions on *m* must again match the previous conditions on *abm*, since *a*; *b* > 0. With an analogous reasoning for 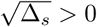, we ultimately obtain:

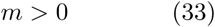

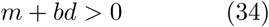

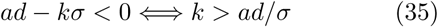

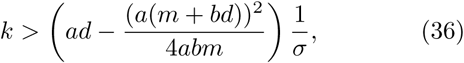

or alternatively:

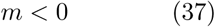

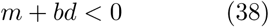

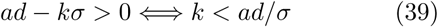

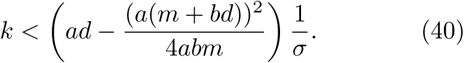

The case iii) is of particular interest, since it entails that there could exist two non-negative cancer attractors (stable fixed points of (1-2)), which are separated by one unstable state. For both solutions 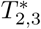 to be positive, we require that 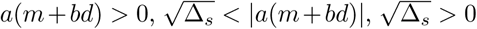 and *abm* > 0, or equivalently, 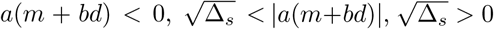 and *abm* < 0. Thus, this is identical to case i), except for the fact the conditions on *a*(*m* + *bd*) are exactly inverted. We can therefore simply adopt the conclusions from the discussions of 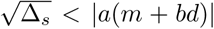 and 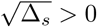.

As in i), the *m* > 0 sub-case of the two possible sub-cases in the discussion of Δ_*s*_ > 0, to obtain 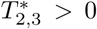, needs to satisfy the additional conditions *a*(*m* + *bd*) > 0 and *abm* > 0. Following the assumptions about the values of *a, b* and *d*, this immediately entails that *m*+*bd* > 0 and that *m* > 0. The other sub-case (*m* < 0) requires *a*(*m* + *bd*) < 0 and *abm* < 0, which by analogy entails that *m* + *bd* < 0.

Hence, summarizing, for all conditions for 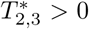 to be satisfied at the same time, there exist two ways in which this can be achieved, that crucially depend on the sign of the effector cell growth stimuation paramter *m*. On the one hand, we will obtain two positive solutions to (1), if:

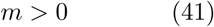

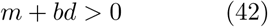

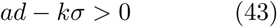

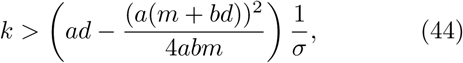

or otherwise, if:

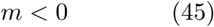

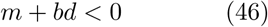

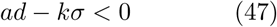

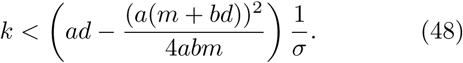

This last set of conditions with *m* < 0 is of particular interest. On the one hand, *m* > 0 leads to biologically dissatisfactory scenarios: The root at 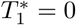 is repulsive, an there exists only one positive attractive root 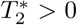. This behavior signifies a departure from the concept of an *immunological barrier* [2, 8], whereby a tumor has to first surpass a threshold size, from which it is hindered by the immune system, before being able to grow to large numbers. On the other hand, the resulting bistability pattern in the *m* < 0 case is reminiscent of key features of cancer establishment and growth, which makes them worth studying.

### Stability analysis

To analyze the stability of the equlibria of the system of equations (1 - 2), we perform a classical analysis based on the behavior of the Jacobian of the map *f*: (*E, T*) → (*dE/dt, dT/dt*) at equilibrium points. To do this, we first interpret (1 - 2) as a map:

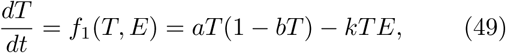

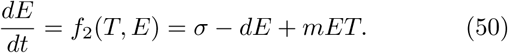

The Jacobian, *J*, of the map *f* is thus:

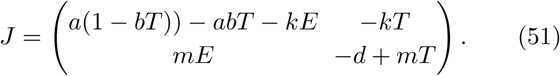

For an equilibrium point to be stable, the trace of *J, tr*(*J*) needs to be negative, while the determinant of *J, det*(*J*), needs to positive. The trace and determinants of *J* must therefore satisfy:

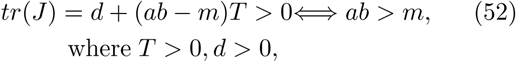

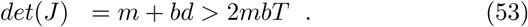

The second condition (53) is equivalent to 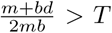 if *m* > 0, and to 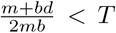 if *m* < 0. If *m* > 0, the largest tumor fixed point is 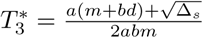, from (3). Inserting 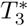 into (53) shows that the condition cannot be satisfied. However, the 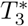 does satisfy *det*(*J*) < 0 if *m* < 0. Thus, only *m* < 0 scenarios can lead to the largest of the fixed points being stable.

## APPENDIX B: MATHEMATICAL ANALYSIS OF THE FREQUENCY DEPENDENT IMMUNE RESPONSE MODEL

To investigate the steady state solutions for the equations (9-10), we first solve for the steady state of *E*, which can be found analytically:

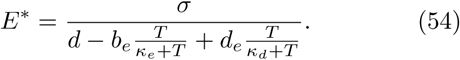

Substituting *E* in the expression for *dT/dt* with equation (54) reduces the finding of the steady states to a one-dimensional problem in *T*\*:

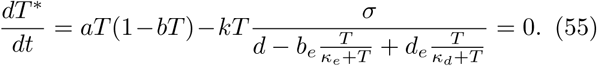

*T*\* = 0 is a trivial root of 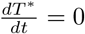. After removing the trivial root from (55), the remaining fixed points of the system (9-10) correspond to the roots of:

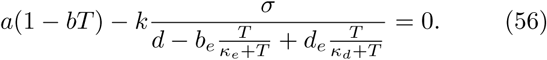

This expression can be converted into a polynomial of third order in *T* *. Solving equation (56) is thus equivalent to the problem of finding the roots of a cubic equation that is obtained from (56) by eliminating the denominators:

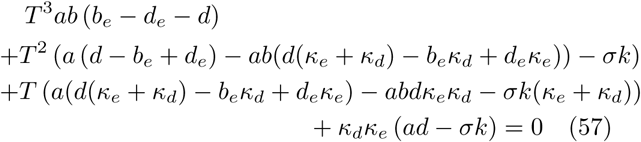

In other words, (57) is equivalent to a cubic equation of the form:

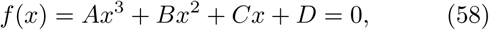

where *T* = *x* and

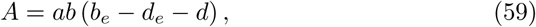

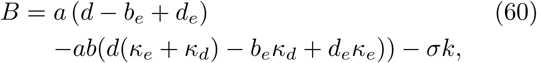

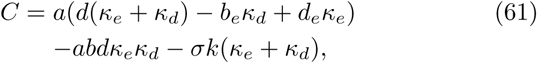

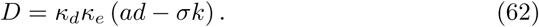

The cubic is analytically solvable, but tedious to write out in its standard representation. We thus employ a non-standard representation of these roots to better analyze the systems’ behavior and its biological meaning. The properties of the roots of (59) are largely dependent on the discriminant Δ of the cubic equation, which is defined by:

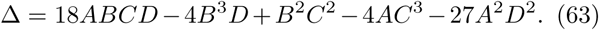

If Δ > 0 all three roots are real and distinct, if Δ = 0 one root is a multiple root, and if Δ < 0 only one root is real, and the other two are complex.

### Appendix B1: Two positive roots of the cubic

With this information, we can make some assertions about the conditions that (58) and Δ must satisfy for bistability to exist. Under bistability, two of the three roots of (57) must be positive. Then, together with the root *T* = 0, there will be three possible roots to produce a bistability effect. The case in which all three are positive will be dicussed next.

Two positive roots of (57) can be attained in two ways: a) Δ = 0, with both the double and the single root being positive and b) Δ > 0 and only the smallest root being negative, with the other two roots both being positive.

The case a) is likely biologically irrelevant, since all the coefficients in (58) would have to take values to exactly satisfy Δ = 0. In case b) where Δ > 0 and all roots exist, we want to find conditions under which exactly two of the three of the roots are positive. To this end, we draw on the theory of cubic functions and on some of its fundamental results about the nature of roots of cubic functions.

An elegant way to represent the roots of the cubic function is by a trigonometric approach. This approach consists in modifying the standard cubic equation by means of the *Tschirnhaus transformation*, where *x* = *t* − *B/*3*A*, and *t* is the new variable. The transformation uses the insight that the sum of the roots *x*_*N*_ of any n-order polynomial *a*_*n*_*x*^*n*^ + *a*_*n*−1_*x*^*n*−1^ … *a*_1_*x*+ *a*_0_ is given by − *a*_*n*-1_*/na*_*n*_. Thus, the Tschirnhaus transformation corresponds to a shift along the x-axis that relocates the cubic such that the sum of the roots lies at the origin [86]. Expressing the standard cubic representation (58) in terms of the variable *t* leads to the *depressed cubic*:

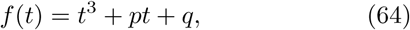

where

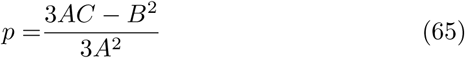

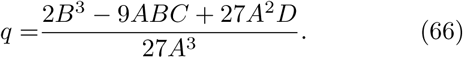

If three real solutions exist (Δ > 0), the roots of the depressed equation can be expressed by the use of trigonometric functions [86, 87]. Then, the real roots of the depressed cubic equation are [88]:

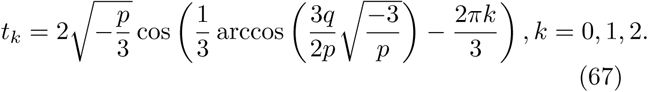

This equation is only valid if the the argument of the arccosine of 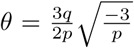 is ∈ (−1, 1). Interestingly, this is equivalent to demanding that 4*p*^3^ + 27*q*^2^ ≤ 0. This condition implies that *p* < 0, also a necessary condition for 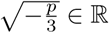. The solution (67) has an elegant geometric interpretation. The roots can be represented as the projections of the vertices of an equilateral triangle with a circle of radius 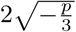 onto the x-axis. The center of the circle is located at *x*_*N*_.

Thus, what is needed to ensure that one real root is negative and two real roots are positive is to demand that the back-transformation of the smallest root min_*k*_ *t*_*k*_ − *B/*3*A* be smaller than zero, and the next largest back-transformed root be positive. Cubic function theory has produced useful results on how to find the smallest solution min_*k*_ *t*_*k*_ [86, 88] as well as the position of the other roots in relation to it. Let *t*_0_ be interpreted as a function of *p* and *q*, that is, *t*_0_ = *C*(*p, q*). Then the three roots can be expressed in terms of *C*(*p, q*), namely

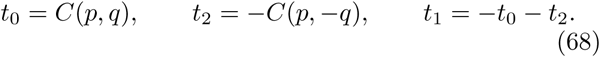

And furthermore, if the three roots are real, we have *t*_0_ ≥ *t*_1_ ≥ *t*_2_ [86, 88]. Thus, *t*_2_ is the smallest of the solutions, and therefore:

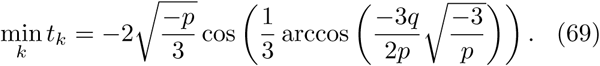

With this, the conditions to obtain one negative and two positive roots can be specified. First, Δ > 0 for all roots to be real-valued, second *t*_2_ − *B/*3*A <* 0 to obtain a negative minimal root, and third, *t*_1_ − *B/*3*A >* 0 to ensure that the second largest root is positive. When replacing the values of *p* and *q* in (64) with terms of the coefficients of the standard cubic function (58), these conditions read as follows:

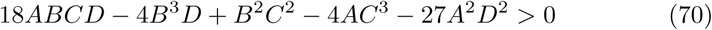

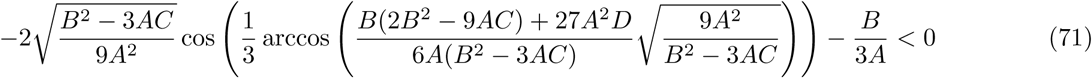

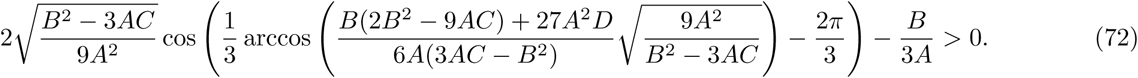

Expressing *A, B, C* and *D* in terms of the parameters of model (9-10) gives the full solutions to the conditions for bistability. These conditions only ensure a certain position of the roots, but do not establish whether these roots are ordered into stable and unstable equilibria in a biologically reasonable way.

### Appendix B2: Three positive roots of the cubic

The trigonometric interpretation of the cubic also allows us to identify the conditions for all roots to be positive. Applying the same reasoning as in the two-positive-roots case (*Appendix B1*), we impose that the sum of the smallest root *t*_*k*_ and the shift implied in the Tschirnhaus transformation, *x*_*N*_ = −*B/*3*A*, is larger than zero. In other words, we ask that the solution *t*_*k*_, which is located the farthest from *x*_*N*_ in negative-x direction should be smaller than *x*_*N*_ itself. In this case, all roots will come to lie on the positive part of the x-axis, although the circle of radius 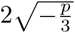 around *x*_*N*_ might reach into negative x-values.

The condition 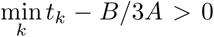 is thus the precise condition for which all roots are positive. By virtue of (69), and inserting the values of *p* and *q* in terms of the coefficients of the standard cubic function (58), the conditions for only positive roots become:

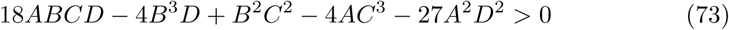

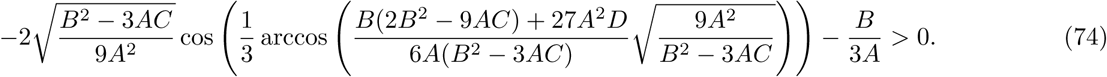

These are implicit inequalities in A, B, C and D, which in turn depend on the parameters of model (9-10) by means of the relation (59).

Expressing the coefficients A, B, C and D in terms of the parameters of model (9-10) thus gives the exact analytical conditions under which four non-negative fixed points to (9-10) exist, out of which three are strictly positive. Writing this expression down is cumbersome and hard to analyze further. However, numerical methods can be employed to examine which regions in parameter space can produce this distinctive pattern of multistability.

### Appendix B3: Analyzing parameter space for existence of multistability

We analyzed whether the conditions (70) and (73) are satisfied for realistic parametrizations of model (9-10). To this end, we fixed most parameters at biologically plausible values found in the literature (see Table II). We then varied two of those parameters which are either expected to have a large impact on the system’s dynamics, or may be less well understood.

The standard parameter values used for the extended model with saturation are specified in Table II.

Figure 5 shows the regions of parameter space whose parameter values lead to bistability as well as multistability in the model (9-10). The region that supports multistability forms a band across a wide range of immune cell exhaustion rate values *d*_*e*_. This indicates that when a waning of killing efficacies *k* occurs, the system may trespass this region. A boundary is shared by a region with bistability and the parameter region that only admits a cancer-free state. The mechanism for combination of immunotherapy may thus still be applicable along this boundary, which is present at plausible values of *d*_*e*_ around unity. The rest of the parameter values used for the generation of this plot are within the range of possible values frequently used in cancer modeling.

**FIG. 5.**
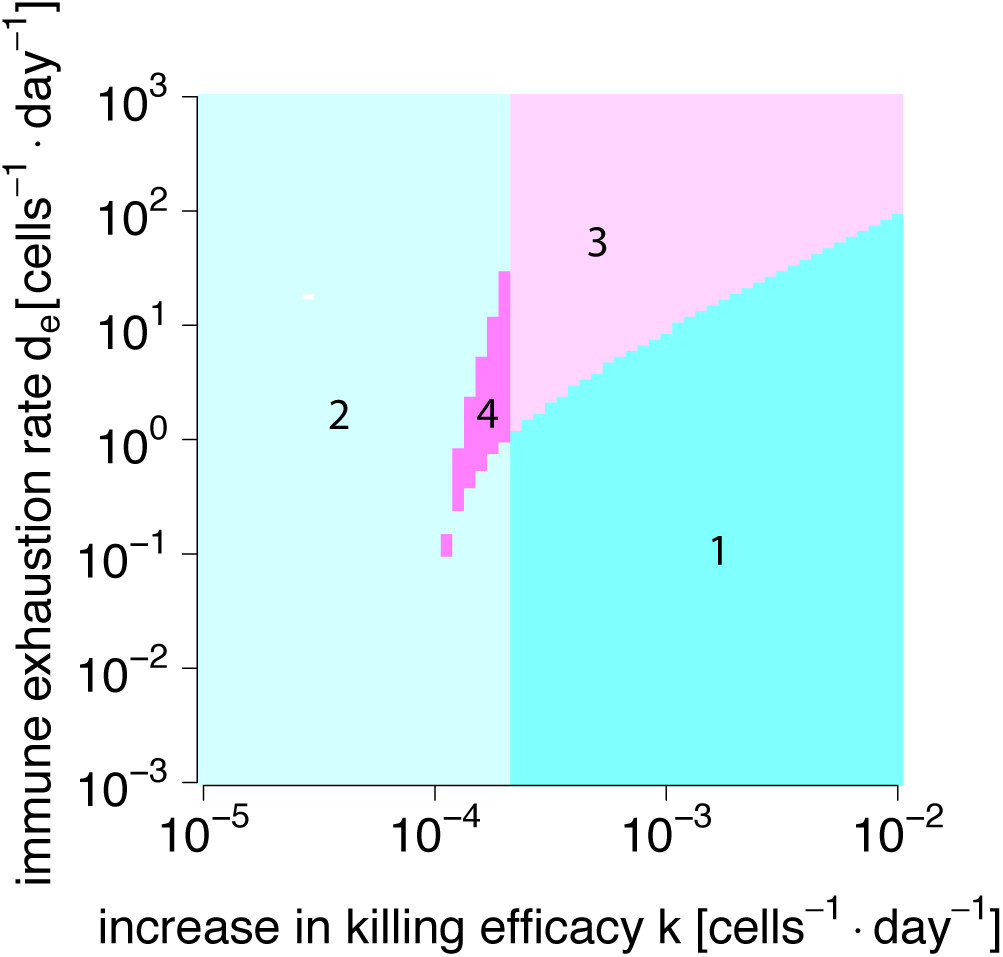
Number of possible non-negative steady states of *T* of the model (9-10). Shown is the region of parameter space spanned by killing efficacy *k* and effector cell exhaustion *d*_*e*_ separated according to the non-negative fixed points of the model (9-10). The number of roots were identified numerically by means of the *rootSolve* package in R [50–52]. Numbers indicate the number of non-negative fixed points in each parameter region. The region that supports multistability is colored in violet. The region that admit bistability (3 fixed points) is adjacent to the multistability region. The other used paramaters are as in Table II.

**FIG. 6.**
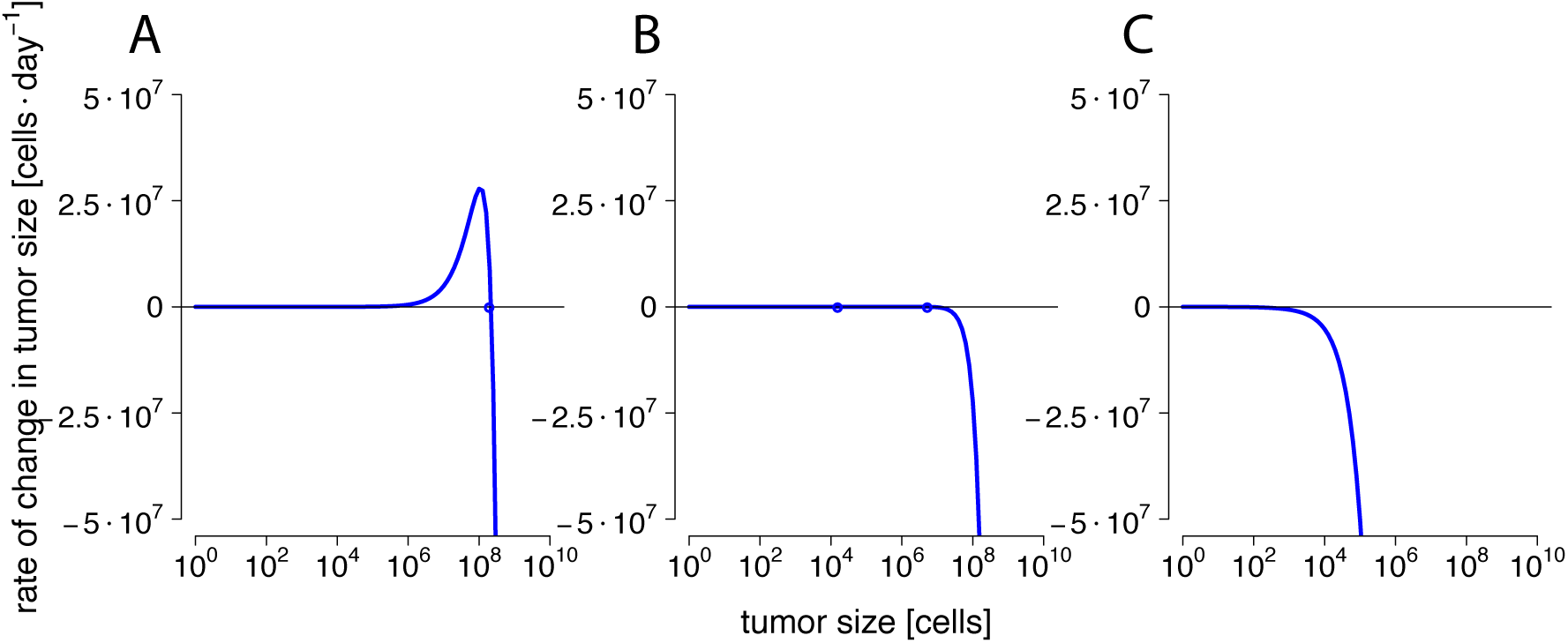
Bistability of cancer cell fixed points induced by immune control of tumor growth in model (20-22). Panels A) to C) show the function *dT/dt*(*T*) (blue lines) for different parameter values of *c. c* is varied from 10^−7^ (A) to 10^−3^ (B) 10^0^ (C). The blue points indicate the roots of *dT/dt*(*T*). Bistability emerges at *c* = 10^−3^. The parameter values were as specified in Table III.

**TABLE III.** Parameter values for the extended model with saturation in NK and CTL (20-22).

| Parameter | Description | Value |
| --- | --- | --- |
| $a$ | maximum replication rate of unmutated tumor cells [5] | $0.514 \text{ day}^{-1}$ |
| $b$ | inverse carrying capacity of tumor [5] | $1.02 \cdot 10^{-9} \text{ cells}^{-1}$ |
| $c$ | killing efficacy natural killer cells | $10^{-7} - 10^0 \text{ cells}^{-1} \text{ day}^{-1}$ |
| $k$ | killing efficiency of effector cells or CTLs [41, 57, 58] | $10^{-5} - 10 \text{ cells}^{-1} \text{ day}^{-1}$ |
| $\sigma$ | replenishment rate of effector cells or CTLs $E$ [34, 35] | $10 - 10^3 \text{ cells} \cdot \text{day}^{-1}$ |
| $b_n$ | proliferation rate of natural killer cells $N$ | $1.5 \text{ cells}^{-1} \text{ day}^{-1}$ |
| $d_n$ | exhaustion rate of natural killer cells $N$ | $1 \text{ cells}^{-1} \text{ day}^{-1}$ |
| $\mu$ | death rate of natural killer cells $N$ | $1 \text{ day}^{-1}$ |
| $d$ | effector loss rate [36] | $2 \text{ day}^{-1}$ |
| $d_e$ | CTL immune impairment rate [36, 67] | $2 \text{ day}^{-1}$ |
| $b_e$ | maximum CTL proliferation rate [36, 66] | $1 \text{ day}^{-1}$ |
| $\omega$ | growth stimulation rate of cytotoxic T lymphocytes $E$ by natural killer cells [5] | $1.1 \cdot 10^{-7} \text{ cells}^{-1} \text{ day}^{-1}$ |
| $\kappa_{be}$ | saturation coefficient of CTL immune growth | $5 \cdot 10^2 \text{ cells}$ |
| $\kappa_{de}$ | saturation coefficient of CTL immune exhaustion | $2.5 \cdot 10^4 \text{ cells}$ |
| $\kappa_{bn}$ | saturation coefficient of NK immune growth | $10^4 \text{ cells}$ |
| $\kappa_{dn}$ | saturation coefficient of NK immune exhaustion | $10^3 \text{ cells}$ |

## APPENDIX C: MATHEMATICAL ANALYSIS OF NK-CTL MODEL

We follow the approach taken for both the base model and the extended base model with saturation and attempt to derive a function 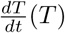 to analyze the roots of (20-22), which are the fixed points of the system. The steady states of *N* and *E* can be found analytically:

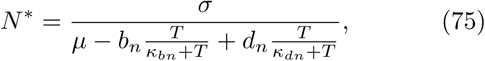

whereas

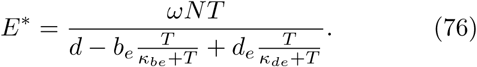

Substituting *N* and *E* in the expression for 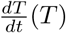 in (20-22) reveals that 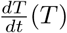 can be expressed as

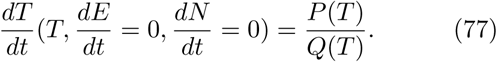

Here, *P*(*T*) = *F*(*T*)*T* is a sixth-order polynomial in *T*. Again removing the trivial solution 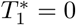, leaves a polynomial *F*(*T*) of fifth order. *Q*(*T*) is a fourth order polynomial. The roots of *P*(*T*) constitute the remaining fixed points of the system. In general, an analytical expression for the roots for such a polynomial can only be given up to the fourth order. Thus, we first restrict ourselves to discussing more general properties that *P*(*T*) and *Q*(*T*) need to satisfy in order to generate biologically plausible situations.

For this, it is important to keep in mind that 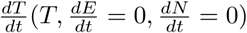 has a narrow mathematical interpretation. In the three-dimensional state space spanned by *T, E* and *N*, it describes the behavior of 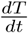 along the intersection between the nullclines of *E* and *N*. In a three-dimensional space, this intersection is typically a line passing through a steady state. Imagine a point with some *T* -value approaching a steady state at *T*\* on the intersecting line. For the steady state to be stable, 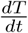 would have to be positive as the point, and with it *T* < *T*\*, reaches *T*\* from below. Equally, as the point moves past the steady state into larger values *T* > *T*\*, 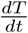 would have to grow negative. We call this a *stabilizing* behavior of 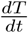 at the steady state. If a steady state satisfies this condition, we call it *T* -stabilizing. Clearly, this behavior of 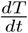 around the steady state does not guarantee that it is stable: instability could still be caused by other properties of the derivatives field perpendicular to the null cline analyzed. However, it is a pre-requisite for the steady state to be stable.

We begin our discussion by analyzing the simpler problem 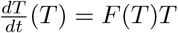. Here, 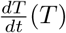 is a polynomial *F*(*T*) times *T*. The order of the polynomial *F*(*T*) and the sign of the leading coefficient *A* of that polynomial, influences the locations of the roots of *F*(*T*), and thus also of 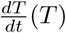. *F*(*T*) also influences how the roots are arranged into stable fixed points.

The *n*-th order polynomial will maximally have *n* real valued roots. If *n* is odd, and *A* < 0, the smallest and largest fixed points of *F*(*T*) are *T* -stabilizing. Conversely, if *n* is odd, and *A* > 0, the smallest and largest fixed points of *F*(*T*) are unstable. If *n* is even, and *A* < 0, the smallest fixed point of *F*(*T*) is unstable, while the largest is *T* -stabilizing. Analogously, if *n* is even, and *A* > 0, the smallest fixed point of *F*(*T*) is *T* -stabilizing, and the largest is unstable.

These claims follow from the intermediate value theorem. If *n* is odd, the limits of the polynomial for *T* → ∞ and *T* → −∞ will be of opposite signs. If the leading coefficient *A* of the polynomial is positive, 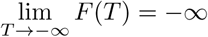, and 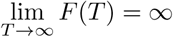. To attain values from −∞ to ∞, *F*(*T*) must traverse the *T* -axis from negative to positive by virtue of the intermediate value theorem. Thus, the first root must be an unstable fixed point. Since there exist an odd number of roots, and the roots must alternate from *T* -stabilizing to unstable, the last root must also be an unstable equilibrium. The converse is true for *A* < 0, with both, the smallest and largest fixed points being *T* -stabilizing. For even values of *n*, if *A* > 0, *F*(*T*) will either be ∞ in the limit *T* → ±∞ and −∞ for *A* < 0. Thus, following the same logic as with *n* odd, the smallest fixed point must be *T* -stabilizing if *A* > 0, while the largest will be unstable. Conversely, if *A* < 0, the smallest fixed point will unstable, whereas the largest will be *T* -stabilizing.

Multiplying *F*(*T*) with *T* does not alter the position of *F*(*T*)’s roots. However, it will have the effect of adding a new root *T* = 0 to 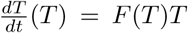, and to switch *T* -stabilizing into unstable (and vice versa) fixed points if they are negative.

With this in mind, we can now turn our attention to the full expression, 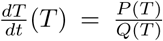. Division by *Q*(*T*) does not alter the position of the roots (unless the roots of *Q*(*T*) are identical with the roots of *P*(*T*), in which case they would cancel out). However, it can modify the *T* -stabilizing properties of the roots. Roots will be *T* -stabilizing if the *T* -derivative of 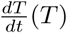 is negative. Let us assume that *F*(*T*) has *n* roots *T*_*i*_ (*i* ∈ {1, …, *n*}). Then,

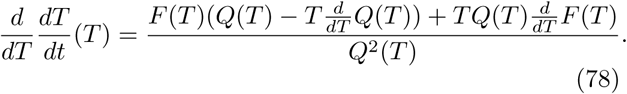

If we evaluate this expression at a stable root of *F*(*T*), *T*_*i*_, where 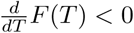, we obtain that:

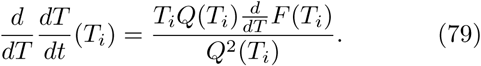

This entails that the a root *T*_*i*_ > 0 (the only biologically plausible solution) will only be stable if *Q*(*T*_*i*_) > 0. Thus, for the roots of *F*(*T*) to retain their *T* -stabilizing properties, they will need to come to lie in intervals of *T* where *Q*(*T*) is positive.

Since the fixed points of (20-22) can be described as *F*(*T*)*T*, the above reasoning is valid, and we can draw conclusions about what conditions the leading coefficient *A* must satisfy to ensure biologically plausible equilibria. Analyzing the structure of 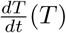 with *Mathematica* [49], reveals that the coefficient *A* = *a*_5_ of the polynomial *F*(*T*) = *a*_5_*T* ^5^ + *a*_4_*T* ^4^ + *a*_3_*T* ^3^ + *a*_2_*T* ^2^ + *a*_1_*T* ^1^ + *a*_0_ reads as follows:

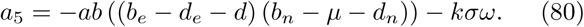

Given that the order of the polynomial *F*(*T*) in NK-CTL model (20-22) is odd, the leading coefficient *a*_5_ of the polynomial must be negative and the value of *Q*(*T*) at the largest fixed point positive in order for *F*(*T*) to have a *T* -stabilizing largest positive fixed point. Thus, analogously to the situation in the extended base model with saturation, the biological plausibility of the positioning of the fixed points is compatible with *a*_5_ < 0. Again, *a*_5_ expresses the balance between proliferative and suppressive forces acting on NK as well as CTL cells, and is independent from the saturation coefficients. As in the previous models, we assume that *a, b, d, σ, k* > 0 and it is biologically reasonable also to assume *ω* > 0. Thus, interestingly, *a*_5_ < 0 can be satisfied by increasing *kσω*. But if *kσω* is negligible, *a*_5_ will only be negative if (*b*_*e*_ − *d*_*e*_ − *d*) and (*b*_*n*_ − *µ* − *d*_*n*_) are both negative or both positive. Thus, the NK-CTL model allows for more flexibility to attain biologically plausible scenarios than the two previous models: if the the balance between proliferative and suppressive forces in both NK and CTLs is tipped in favor of exhaustion of the immune cells, biologically interpretable dynamics can emerge. This is analogous behavior to an *m* < 0 situation in the base model. Similarly, if the balance is tipped towards proliferation in both cell types, the arising scenarios are again biologically sound. However, the behavior between both immune cell types must be similar to attain this.

Unfortunately, the above analysis does not reveal whether bi- or multistability can be guaranteed to arise for special combinations of parameters. To show this, we must fall back to numerical methods. In the following, we prove that the system (20) can generate at least bistability patterns of the kind displayed by the base model. We only analyzed the effects of changing *c*, the killing efficacy of NKs rather than CTLs, on the system. We chose *c* because NK have been estimated to have faster cancer suppressing effects than CTLs in [5]. Figure 6 shows the emergence of the bistability pattern for biologically sound parameter choices in the model (20-22). The parameter values are specified in Table III.

